# Distinct gene-selective roles for a network of core promoter factors in *Drosophila* neural stem cell identity

**DOI:** 10.1101/434597

**Authors:** Alexandre Neves, Robert N. Eisenman

## Abstract

The transcriptional mechanisms that allow neural stem cells (NSC) to balance self-renewal with differentiation are not well understood. Employing an in vivo RNAi screen we identify here NSC-TAFs, a subset of nine TATA-binding protein associated factors (TAFs), as NSC identity genes in *Drosophila*. We found that depletion of NSC-TAFs results in decreased NSC clone size, reduced proliferation, defective cell polarity and increased hypersensitivity to cell cycle perturbation, without affecting NSC survival. Integrated gene expression and genomic binding analyses revealed that NSC-TAFs function with both TBP and TRF2, and that NSC-TAF-TBP and NSC-TAF-TRF2 shared target genes encode different subsets of transcription factors and RNA-binding proteins with established or emerging roles in NSC identity and brain development. Taken together, our results demonstrate that core promoter factors are selectively required for NSC identity *in vivo* by promoting cell cycle progression and NSC cell polarity as well as by restraining premature differentiation. Because pathogenic variants in a subset of TAFs have all been linked to human neurological disorders, this work may stimulate and inform future animal models of TAF-linked neurological disorders.

**Author summary:** The brains of many animal species are built with brain stem cells. Having too many brain stem cells can lead to brain tumors whereas too few can lead to birth defects such as microcephaly. A number of next generation sequencing studies have implicated proteins referred to as TATA-box-binding protein associated factors (TAFs) in human neurological disorders including microcephaly, but prior to this study, their function in brain development was unknown. Here we use brain stem cells, known as neural stem cells (NSCs), from the fruit fly *Drosophila melanogaster* as a model system to decipher how TAFs control brain stem cell identity. By combining genetics and low-input genomics, we show that TAFs directly control NSC cell division and cell polarity but do not appear to be required for NSC survival. We further show that TAFs accomplish these functions by associating either with their canonical partner TBP (TATA-binding protein) or the related protein TRF2. In summary, our study reveals unexpected and gene-selective functions of a unique subset of TAFs and their binding partners, which could inform future studies that seek to model human neurological disorders associated with TAFs.

## Introduction

Brains from a wide range of animal species are populated by Neural Stem Cells (NSCs). NSCs must balance self-renewal with differentiation, and failure to achieve this balance can lead to neurological disorders. For example, excess self-renewal of NSCs has been linked to brain tumors whereas premature depletion of NSCs is thought to contribute to primary microcephaly, a human neurological disorder characterized by markedly reduced brain size at birth (1–3). In the last decade, *Drosophila* NSCs have emerged as a powerful in vivo system to study NSC identity because they provide a genetically tractable model that recapitulates many important features of mammalian NSC biology including asymmetric cell division, coupling between the cell cycle and cell fate and a dependence on conserved polarity complexes (4).

Two major types of NSCs have been shown to populate the *Drosophila* central brain. Type I NSCs express two related bHLH transcription factors, Asense (Ase) and Deadpan (Dpn), and divide asymmetrically into a single type I NSC and into a differentiating daughter, the ganglion mother cell (GMC). The GMC in turn undergoes a terminal division to give rise to two post-mitotic neurons. In contrast, type II NSCs express Dpn but not Ase, and divide into another type II NSC and an Intermediate Neuronal Progenitor (INP). After maturation, the INP expresses Ase and undergoes limited rounds of self-renewing divisions while also generating GMCs, much like a type I NSC (Fig 1A). Asymmetric distribution of cell fate determinants such as the transcription factor Prospero and the RNA-binding protein Brat is thought to control differentiation in type I and type II lineages respectively (5–8). In contrast to the many genes that have been identified to be required for the differentiation of both type I and type II lineages, the gene regulatory networks that control NSC identity are less well understood. Here we define NSC identity as the suite of stem cell attributes such as asymmetric cell division, self-renewal, survival, growth and proliferation that collectively distinguish NSCs from their differentiating and differentiated progeny.

**Figure 1.**
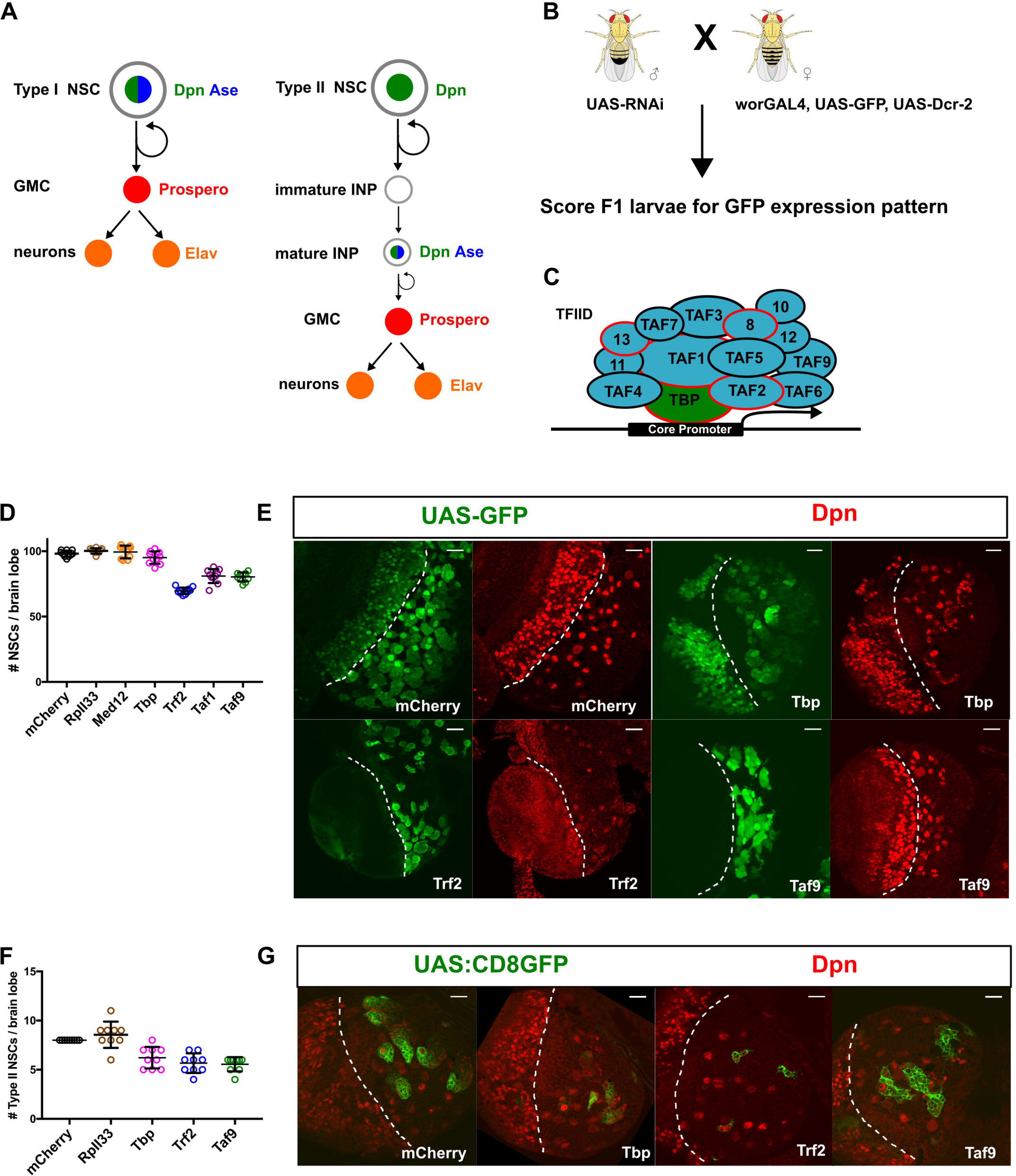
A unique subset of TAFs (NSC-TAFs) required for NSC homeostasis. (A) Marker gene expression and cell division patterns of Type I (left) and Type II (right) NSCs. (B) Genetic schema of *in vivo* transgenic RNAi screen. (C) Subunit composition of Transcription Factor IID (TFIID). Subunits with a red outline have been implicated in human neurological disorders. (D) Quantification of total Central Brain NSCs (large Dpn^+^ cells) in nervous systems expressing the listed RNAi transgenes using a pan-NSC driver (worniuGAL4, UAS-Dcr-2, UAS-GFP). p values were RpII33 (0.6655), MED12 (0.9376), TBP (0.3182), TRF2 (<0.001), TAF1 (<0.001), TAF9 (<0.001). (E) Single confocal section of brain lobes expressing the indicated RNAi transgenes showing expression of either UAS-GFP (left) or the NSC marker Dpn (right). (F) Quantification of Type II NSCs in nervous systems expressing the listed transgenes using a Type II NSC driver (UAS-Dcr-2; worniuGAL4, aseGAL80, UAS-CD8:GFP). CD8:GFP is a membrane targeted GFP transgene. p values were RpII33 (0.5449), TBP (0.0011), TRF2 (<0.0001), TAF9 (<0.0001). (G) Single confocal section of brain lobes expressing the listed RNAi transgenes showing expression of either UAS-CD8:GFP (left) or the NSC marker Dpn (right). In D and F, experimental genotypes were compared to the mCherry control using a one-way ANOVA with Dunnett’s multiple comparison test. In E and G, the white dotted line demarcates the optic lobe (left)/ central brain (right) boundary and scale bars represent 20 μm. Note that TRF2 or TAF9 depletion leads to increased GFP expression in the central brain whereas TBP depletion leads to decreased GFP expression; compare with optic lobe GFP expression.

In order to understand in more detail the basis for NSC identity we carried out a focused RNAi screen in live *Drosophila*, testing transcriptional regulators that influence the number of NSCs and the size of NSC lineages (Fig 1B). Unexpectedly we found that a subset of TAFs (TBP-associated factors) and the TBP-related factor 2 (TRF2) are required for maintaining normal NSC numbers, NSC proliferation and NSC cell polarity, but do not appear to be required for NSC survival. We further find that NSCs depleted for TAFs or TRF2 are hypersensitive to cell cycle manipulation. TAFs have been well characterized as subunits of the ~1 megadalton Transcription Factor IID (TFIID) complex comprised of the TATA box-binding protein (TBP) and 13 individual TAFs (Fig 1C(9)). The main function of TFIID is to recognize and bind to the core promoter, a segment of DNA that is sufficient to direct accurate and efficient RNA polymerase II transcription. In addition to contributing to core promoter recognition, individual TAFs are known to have distinct biochemical activities. For example, TAF4 has been shown to function as a coactivator for different transcription factors through its conserved ETO domain (10).

A number of recent studies suggest that some TFIID subunits are neither universally required for gene expression nor invariant (9, 11–13). However, while different subsets of TFIID subunits have been shown to be required for the self-renewal of both murine and human embryonic stem cells (ESCs), the function of TAFs in stem cell populations in vivo has not been investigated (14, 15). Notably, a number of recent genetic studies have identified pathogenic variants in several TFIID subunits. First, variants in *TAF1*, *TAF2, TAF8* and *TAF13* have all been linked to intellectual disability and microcephaly (16–19). Second, insertion of an SVA-type retrotransposon in a noncoding region of *TAF1,* that results in abnormal splicing and reduced expression of *TAF1* in patient-derived NSCs, is associated with X-linked dystonia parkinsonism (20, 21). Third, *TBP* is a candidate microcephaly and intellectual disability gene in patients with a subtelomeric 6q deletion, whereas *de novo* expansion of CAG repeats in *TBP* is thought to cause spinocerebellar ataxia 17 (22, 23). Finally, mutations in *TAF6* have been linked to a Cornelia de Lange-like syndrome, a clinically heterogeneous disorder characterized by developmental delay and intellectual disability (24). In this study, we extend the role of TAFs, TBP and TRF2 in developmental gene regulation by showing they are members of a core promoter network involved in NSC identity.

## Results

### A subset of TAFs is required for NSC identity

To identify genes that regulate neural stem cell (NSC) identity, we performed a focused transgenic in vivo RNAi screen using an NSC-enriched GAL4 driver line (worniu-GAL4, UAS-GFP, UAS-Dcr-2) to drive expression of either long or short hairpin RNAs (Fig1B). We used the intensity and pattern of the GFP signal in live wandering larvae as a proxy for NSC identity. After testing more than 900 RNAi transgenes, we focus here on the functional analysis of several hits targeting genes encoding TATA box-binding protein-associated factors (TAFs). We found that knockdown of any one of nine different TAFs (*Taf1*, *Taf2*, *Taf4*, *Taf5*, *Taf6*, *Taf7*, *Taf8*, *e(y)1/Taf9*, *Taf12*) resulted in a complex suite of similar phenotypes: fewer NSCs, with remaining NSCs clustering together, smaller NSC lineages, abnormal morphology, reduced expression of the type I NSC marker Ase whereas expression of the pan-NSC marker Dpn remained unaffected (Fig 1D, E; see Fig 2 D,F,H for Ase expression). While the observed NSC loss was moderate, it was highly reproducible and comparable to that observed upon either knockdown or loss of function of other NSC identity genes (25–27). To test whether increased levels of NSC-TAFs can promote NSC fate, we overexpressed a number of different NSC-TAFs as well as additional core promoter factors but did not observe any overt phenotypes (Fig S1A, B). As a positive control, we overexpressed a constitutively active form of aPKC (28), a known NSC identity gene and observed the expected increase in brain size. Intriguingly, knockdown of other prototypical TAFs (*bip2/Taf3*, *Taf10*, *Taf10b*, *Taf11*, *Taf13*) did not result in overt phenotypes (Figure S2 A, B, Table S1). The lack of an observable NSC phenotype for these TAFs is unlikely to be due to inefficient RNAi since knockdown of either *Taf3* or *Taf13* with actin-GAL4 resulted in highly penetrant organismal lethality, findings that are consistent with an independent genome-wide RNAi screen (29). Moreover, NSC clones that were homozygous for lethal alleles of *Taf11* (*Taf11*^*M1*^ and *Taf11*^*M5*^)(30) or *Taf13* (*Taf13*^*LL04552*^)*(31)*, generated using the MARCM (**M**osaic **A**nalysis with a **R**epressible **C**ell **M**arker)(32) system, were either modestly reduced in size (*Taf11*^*M1*^ and *Taf1^M5^*) or indistinguishable (*Taf13*^*LL04552*^) from their wild-type counterparts both in terms of NSC clone size and Ase expression (Fig S2 A, C).

**Figure 2.**
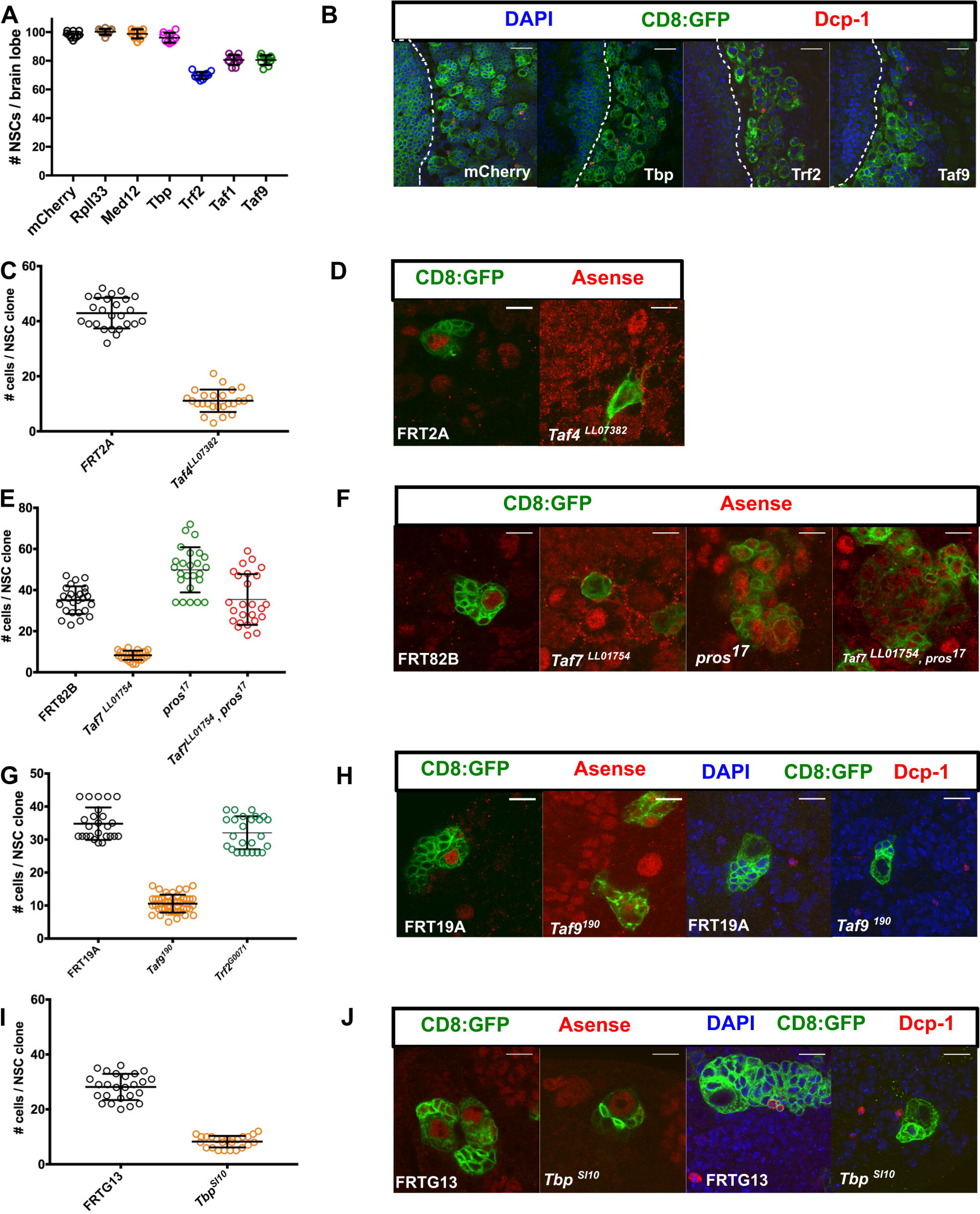
TBP, TRF2 and NSC-TAFs are required for normal NSC output but not for NSC survival. (A) Blocking apoptosis does not rescue NSC loss observed upon knockdown of TRF2, TAF1 or TAF9. The listed RNAi transgenes were co-expressed with UAS-miRHG, which blocks apoptosis, using a pan-NSC driver (worniuGAL4, UAS-GFP). p values were RpII33 (0.4216), MED12 (0.9893), TBP (0.4710), TRF2 (<0.001), TAF1 (<0.001), TAF9 (<0.001). (B) Knockdown of TBP, TRF2 or TAF9 does not result in cleaved caspase (Dcp-1) The white dotted line demarcates the optic lobe (left)/ central brain (right) boundary. (C, D) *Taf4*^*LL07382*^ NSCs generate fewer cells (p value < 0.0001) and exhibit decreased Asense expression. (E, F) *Taf7*^*LL07382*^ NSCs exhibit a marked decrease in Asense expression, generate fewer cells but can be partially rescued by deleting *prospero*. p values were *Taf7*^*LL07382*^ (< 0.0001), *pros*^*17*^ (< 0.0001), *Taf7*^*LL07382*^, *pros*^*17*^ (0.9986) (G, H) *Taf9*^*190*^ NSCs, but not *Trf2*^*G0071*^ NSCs, exhibit a marked decrease in Asense expression and generate fewer cells. p values were *Taf9*^*190*^ (< 0.0001), *Trf2*^*G0071*^ (0.0331). (I, J) *Tbp*^*SI10*^ NSC generate fewer cells but maintain Asense expression (p value < 0.0001). In C and I, experimental clones were compared to the control clones using an unpaired t-test. In A, E and G, experimental clones were compared to control clones using a one-way ANOVA with Dunnett’s multiple comparison test. Scale bars represent 20 μm in B and 10 μm in D, F, H and J.

### TBP, TRF2 and TAF depletion results in shared and private phenotypes

For clarity, we refer to the subset of TAFs that presented a phenotype in our screen as NSC-TAFs. *Drosophila* TAFs are known to be present in either Transcription Factor IID (Fig 1C; TFIID, composed of TBP and 13 TAFs) or SAGA (Spt-Ada-Gcn5-acetyltransferase; including TAF9, TAF10b, TAF12). However, knocking down TBP or SAGA-specific subunits (Gcn5, Spt3, Saf6, Spt7) did not cause a reduction in NSC numbers, or affect Ase expression, despite a marked reduction in the both the GFP signal and organismal viability (Fig 1D; Table S1). Moreover, reducing expression of other components of the transcription initiation complex by knocking down subunits of RNA Polymerase II (RpII33) or the Mediator (MED12) complex, reduced the size of NSC lineages without decreasing NSC numbers (Fig 1D). These data strongly suggest that gene-selective functions of TAFs, rather than general transcriptional activity, are required for NSC identity.

Metazoan lineages evolved several paralogs of TBP called TBP-related factors (TRFs), with TRF1 and TRF3 only found in insect and vertebrate clades, respectively, whereas TRF2 is conserved in all bilaterians (33). We were intrigued that TBP knockdown and NSC-TAF knockdown phenotypes were different, since TAFs and TBP are thought to be obligate partners. We hypothesized that in addition to functioning with TBP as part of TFIID, TAFs may also function with a TBP paralog. Based on previous studies (34–37), the most likely candidate is TRF2. Indeed, knockdown of TRF2, but not TRF1, in NSCs also resulted in reduced NSC numbers, smaller NSC lineages and abnormal NSC morphology (Fig 1D, E, Table S1). Because NSC-TAFs and TRF2 regulate the expression of the type I NSC marker Ase, we asked whether they are also important in type II NSCs, which do not express Ase. Knockdown of either TBP, TRF2 or TAF9 using a type II NSC driver (UAS-Dicer2; worniuGAL4, aseGAL80; UAS-CD8:GFP) diminished the number of type II NSCs (Fig1 F, G). Altogether, our genetic analysis using in vivo transgenic RNAi demonstrate that a unique subset of TAFs, TBP and TRF2 have distinct and gene-selective functions in the identity of both type I and type II NSCs.

### Analysis of *Tbp, Trf2 and Taf* alleles

To corroborate our RNAi results, we assessed the effects of available mutant *Taf* alleles on NSC phenotypes. Because *Tafs* are known or predicted to be essential genes, and loss of *Taf* function in other cellular contexts results in either cell death or cell cycle arrest, we turned again to the MARCM system to analyze NSC clones. In these and subsequent experiments, our choice of which NSC-TAF to analyze was primarily governed by reagent availability. NSC clones that were homozygous mutant for either *Taf4^LL07382^ (31)*, *Taf7^LL01754^(31)* or *Taf9*^*190*^ (38) exhibited a marked reduction in clone size, mitotic index and Ase levels (Fig 2 C-H). Next, we analyzed the phenotype of a *Tbp* allele, *Tbp^SI10^,*that generates a truncated form of the protein that is unable to bind DNA *in vitro (23)*. Despite the fact that TBP is thought to be required for transcription by all three RNA polymerases, *Tbp*^*SI10*^ NSC clones were readily recovered, expressed normal levels of Ase, but generated fewer cells than the control clones (Fig 2 I, J). We anticipated that *Trf2* mutants would phenocopy the examined *Taf* mutants, yet NSC clones homozygous for a hypomorphic allele, *Trf2*^*G0071*^, only exhibited a modest reduction in clone size (Fig 2 G).

The reduced clone size in *Taf4*^*LL07382*^, *Taf7*^*LL01754*^ or *Taf9*^*190*^ or *Tbp*^*SI10*^ NSC clones is indicative of either a delay in cell cycle progression or reduced survival. To distinguish between these possibilities, we examined a marker of apoptosis in either NSCs depleted for TBP, TRF2 or TAF9 by RNAi or *Taf4*^*LL07382*^, *Taf7*^*LL01754*^, *Taf9*^*190*^, *Tbp*^*SI10*^ NSC clones, but did not detect apoptotic NSCs as determined by staining for cleaved caspase (Dcp-1; Fig 2 B, H, J) in any of the genotypes examined. To test whether blocking apoptosis affected the observed NSC loss phenotype, we used a microRNA transgene (miRHG) that concurrently downregulates three major pro-apoptotic genes in *Drosophila*: *reaper*, *hid* and *grim (39)*. Expression of miRHG in NSCs failed to suppress the phenotypes caused by knockdown of TRF2, TAF1 or TAF9 or in NSC clones that were homozygous mutant for *Taf4*^*LL07382*^ (Fig 2A, compare with Fig 1D; Fig S2B). Importantly, we found that the function of NSC-TAFs and TRF2 in cell survival was context-dependent, as knockdown of TBP, TRF2 or TAF9 resulted in apoptosis when the respective RNAi transgenes were expressed in epithelial cells of the wing disc using an inducible GAL4 driver (apterous-GAL4, UAS-GFP; tubulinGAL80^ts^; Fig S2 D).

### TBP, TRF2 and NSC-TAFs control NSC proliferation

Because the reduction in NSC lineage size upon NSC-TAF or TRF2 depletion does not appear to be caused by apoptosis, we predicted that NSC-TAF or TRF2 knockdown NSCs would exhibit a cell cycle delay. To directly test this, we performed EdU labeling experiments and found that after a 2hr pulse, RpII33, TBP, TRF2, TAF1 and TAF9 knockdown NSCs had fewer EdU^+^ NSCs (Fig 3A). All genotypes examined also had a lower mitotic index (pH3^+^ NSCs / total NSCs; Fig 3B). Intriguingly, *RpII33* knockdown NSCs presented a more severe proliferation phenotype, suggesting that the severity of cell cycle progression defects don’t always correlate with NSC self-renewal. (Fig 3B). Examination of additional cell cycle markers demonstrated that TAF1 and TAF9 were required for PCNA-GFP expression, whereas only TRF2 was required for high levels of CycE expression (Fig 3E, F). Accumulating evidence from diverse stem cell systems suggest an intricate link between cell cycle progression and stem cell self-renewal. For example, during development of the mouse cerebral cortex, lowering CyclinD1/Cdk4 expression in NSCs lengthens G1 and results in precocious neurogenesis and similarly, the short G1 phase in murine ESCs has been linked to pluripotency (40, 41). Recent studies showed that *Drosophila* NSCs cultured in vitro increase their cell cycle time during larval development although it’s unclear whether this also occurs through altering the length of G1(42). Because NSC-TAF and TRF2 knockdown NSCs exhibited cell cycle delay, we hypothesized that they had an extended G1 phase and would be hypersensitive to manipulation of the G1/S transition. To address this, we lowered CycE/Cdk 2 activity in control or TAF9 knockdown NSCs, by overexpressing the CIP/KIP family Cyclin-dependent kinase inhibitor Dacapo (Dap). We found that a Dap overexpression had no effect on NSC numbers in a control RNAi background, but further reduced the number of NSCs upon TAF9 knockdown (Fig 3C). Next, we asked whether this sensitivity was specific to the G1/S transition or if it also applies to the G2/M transition. To this end, we overexpressed Wee1, a Ser/Thr kinase that inhibits Cdk1, or knocked down string/cdc25, a phosphatase and Cdk1 activator, in either control or TAF9 knockdown NSCs. Again, we found that Wee1 overexpression or string/cdc25 knockdown did not result in NSC loss, despite an almost complete block in cell cycle progression. However, in combination with TAF9 knockdown, Wee1 overexpression or string/cdc25 loss resulted in a more severe NSC loss relative to TAF9 RNAi alone (Fig 3D). Finally, we asked whether TBP, TRF2 or TAF9 knockdown NSCs exhibited a lengthening of the G1 phase of the cell cycle using the Fly-FUCCI system, a dual color fluorescent reporter for cell cycle stages (43). We further distinguished G2 cells (GFP^+^, RFP^+^, pH3^−^) from cells in mitosis (GFP^+^, RFP^+^, pH3^+^) by staining with the mitotic marker pH3. Surprisingly, we found that the majority of both control and experimental NSCs were in G2. (Fig 3G). Taken together, our results show that NSC-TAFs regulate multiple aspects of NSC cell cycle progression and that depletion of either NSC-TAFs or TRF2 renders NSCs hypersensitive to both G1/S and G2/M cell cycle manipulation.

**Figure 3.**
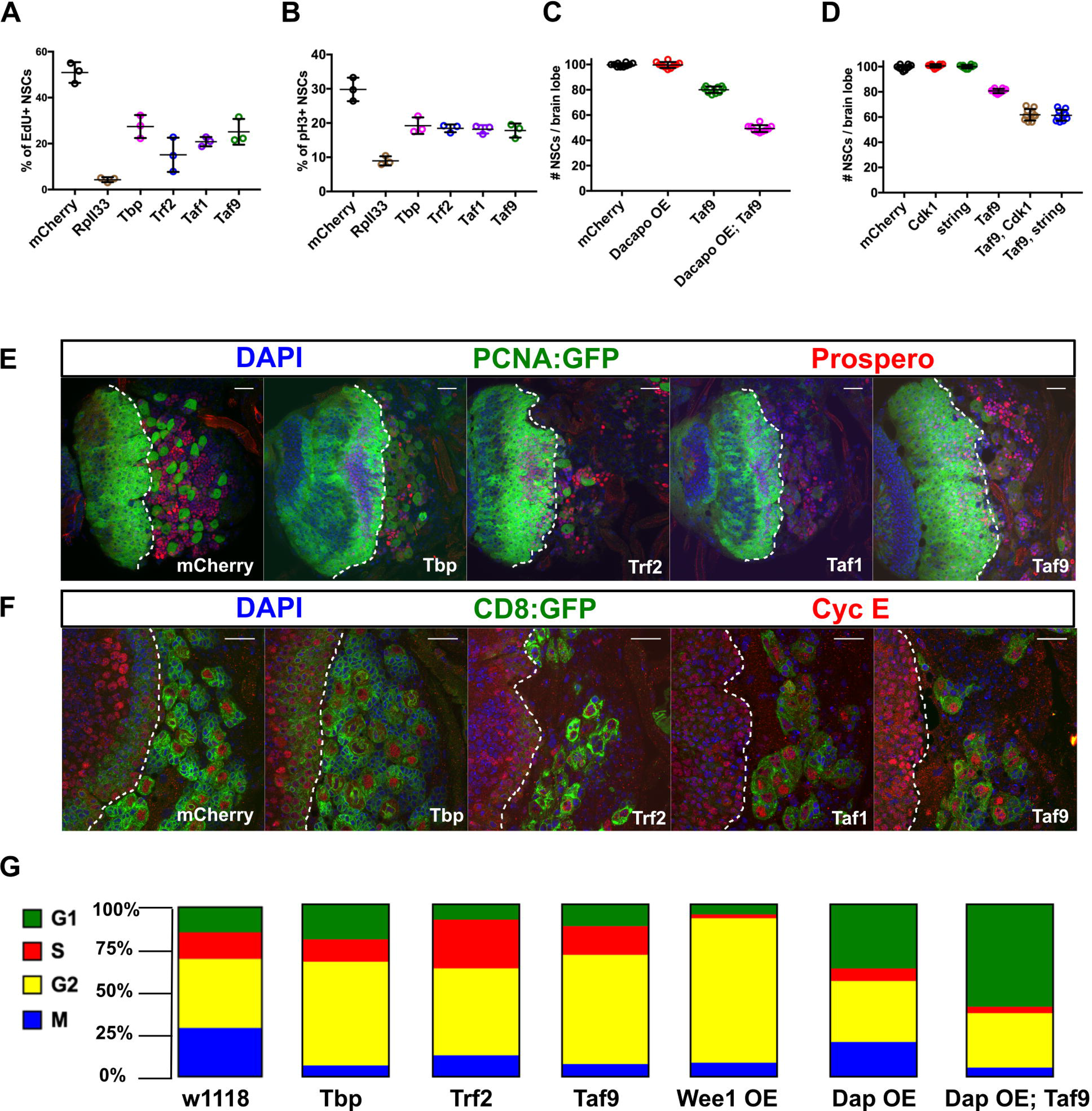
TBP, TRF2 and NSC-TAFs are required for NSC cell cycle progression. (A) Quantification of percentage of NSCs that incorporated the thymidine analog EdU after a 2hr pulse. p values were RpII33 (< 0.0001), TBP (0.0003), TRF2 (< 0.0001), TAF1 (< 0.0001), TAF9 (0.0001). (B) Quantification of percentage of NSCs that are positive for the mitotic marker phospho Histone H3 (pH3). p values were RpII33 (< 0.0001), TBP (0.0002), TRF2 (0.0001), TAF1 (< 0.0001), TAF9 (0.0001). (C) Quantification of NSCs in nervous systems expressing the listed RNAi or overexpression transgenes. p values were Dacapo overexpression (0.9994), TAF9 (< 0.0001), TAF9, Dacapo overexpression (< 0.0001). (D) Quantification of NSCs in nervous systems expressing the listed RNAi transgenes. p values were Cdk1 (0.8166), string (0.9871), TAF9 (< 0.0001), TAF9, Cdk1 (<0.0001), TAF9, string (< 0.0001). (E) Brain lobe images of nervous systems expressing the listed RNAi transgenes, the cell cycle marker PCNA-GFP (Green) and Prospero (Red). (F) Single confocal sections of nervous systems expressing the listed RNAi transgenes and the cell cycle marker Cyclin E. In E and F, the white dotted line demarcates the optic lobe (left)/ central brain (right) boundary (G) Analysis of cell cycle phasing with the Fly-FUCCI system. In A-F, transgenes were expressed with a pan-NSC driver (worniuGAL4, UAS-GFP) and experimental genotypes were compared to the mCherry control using a one-way ANOVA with Dunnett’s multiple comparison test. In G, UAS transgenes were driven by a pan-NSC driver combined with the Fly-FUCCI cassette (worniuGAL4; UASp-GFP.E2f1.1-230, UASp-mRFP1.CycB.1-266/TM6B).

### Regulation of NSC cell polarity and differentiation by TBP, TRF2 and NSC-TAFs

NSC cell polarity is required for asymmetric segregation of cell fate determinants and NSC self-renewal. To address whether the NSC-TAFs and TRF2 phenotypes involve defects in cell polarity, we examined the expression and localization of key polarity proteins that are inherited by the daughter NSC. We found that both interphase and mitotic NSCs that were homozygous mutant for either *Taf4*^*LL07382*^ or *Taf7*^*LL01754*^ failed to express normal levels of Inscuteable (Insc) (Fig 4 A, B). We next examined the localization of atypical protein kinase C (aPKC) in pH3^+^ NSCs. In 23/24 of control NSCs (mCherry), aPKC exhibited a strong cortical crescent. In contrast, only 13/19 of TBP knockdown NSCs, 14/20 TRF2 knockdown NSCs or 5/13 TAF9 knockdown NSCs properly localized aPKC during mitosis (Figure S4B).

**Figure 4.**
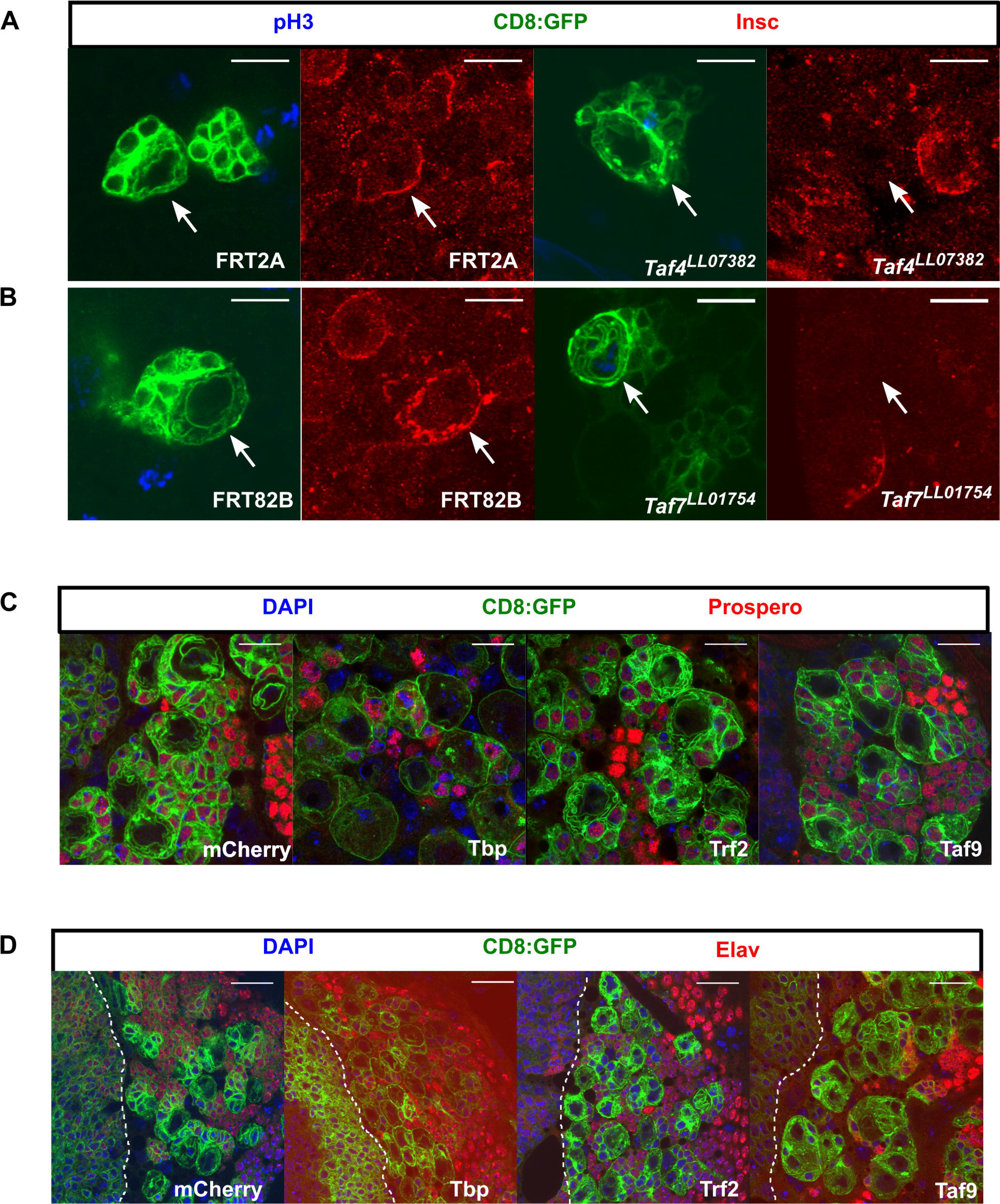
NSC-TAF or TRF2-depleted NSCs exhibit defective cell polarity but do not express differentiation markers. (A) *Taf4*^*LL07382*^ NSCs or (B) *Taf7*^*LL07382*^ NSCs fail to express normal levels of Insc. Mitotic cells are labeled with phospho Histone H3 (pH3), CD8:GFP (a membrane-tethered GFP) labels the clones and Insc is shown in red. (C) Knockdown of TBP, TRF2 or TAF9 does not result in nuclear accumulation of Prospero. (D) Knockdown of TBP, TRF2 or TAF9 does not result in nuclear accumulation of Elav. The white dotted line demarcates the optic lobe (left)/ central brain (right) boundary. In both C and D, transgenes were expressed using a UAS-Dcr2; inscGAL4, UAS-CD8:GFP, tubGAL80^ts^ driver and transgenes induced for 72h. Scale bars represent 10 μm in A, B, C and 20 μm in D.

To ascertain if knockdown of NSC-TAFs or TRF2 results in premature differentiation, we examined the expression of both Prospero (Pros), a homeodomain transcription factor that is necessary and sufficient for differentiation of type I NSCs (5), and Elav, a conserved RNA-binding protein that is expressed in post-mitotic neurons. Neither wild-type NSCs nor NSCs that were depleted for TBP, TRF2 or TAF9 by RNAi or *Taf4*^*LL07382*^, *Taf7*^*LL01754*^, *Taf9*^*190*^ homozygous mutant NSCs, expressed either Pros or Elav in their nuclei (Fig 4C, D, Fig S4C). Moreover, our RNA-seq analysis, described in more detail below, showed that the levels of *pros* and *elav* transcripts were reduced in TAF9-depleted NSCs whereas *pros* but not *elav* trancripts were reduced in either TRF2-depleted or TBP-depleted NSCs. Next, we tested candidate NSC regulators for their ability to suppress NSC-TAF phenotypes and found that overexpression of CyclinD/Cdk4 or a constitutively active form of aPKC (aPKC^CAAX^) did not suppress *Taf4*^*LL07382*^ phenotypes (Fig S2B). However, while Pros was not detected in *Taf7*^*LL01754*^ mutant NSCs, complete loss of *pros* function was able to partially rescue *Taf7*^*LL01754*^ mutant NSC phenotypes (Fig 2E). We conclude that NSCs lacking NSC-TAF or TRF2 exhibit defective NSC cell polarity but unlike other NSC identity genes, do not appear to undergo Pros- or Elav-mediated premature differentiation (26, 27).

### Identification of TBP, TRF2 and NSC-TAF regulated genes

Because the NSC-TAF and TRF2 phenotypes were similar, we hypothesized that NSC-TAFs and TRF2 co-regulate a subset of NSC-expressed genes that in turn are collectively required for NSC identity, and that these NSC-TAF-TRF2 co-regulated genes are distinct from NSC-TAF-TBP-regulated genes. Several lines of evidence suggest that changes in abundance of NSC-TAFs, TBP and TRF2 alone are unlikely to explain their NSC-selective functions. First, using reporters or antibodies, we found that TBP, TAF1, TAF4 and TAF12 proteins are not restricted to NSCs and are expressed ubiquitously in the brain (Fig S3 A-C). We corroborated and extended these observations by mining a published RNA-seq data set, which revealed that TBP, TRF2 and NSC-TAFs are all expressed at comparable levels in FACS-purified NSCs and FACS-purified neurons (44). Second, recent RNA-seq data derived from robotically-sorted NSCs revealed that TRF2 and TAF4 were among a group of 61 transcription factors highly expressed in multiple NSC lineages (45). Therefore, we reasoned that understanding how NSC-TAFs and TRF2 regulate NSC identity requires the identification of NSC-specific transcriptomes. To identify genes co-regulated by NSC-TAFs and TRF2, we performed RNA-seq analysis of FACS-purified NSCs expressing either control (mCherry), TBP, TRF2, or TAF9 RNAi transgenes in type I NSCs, driven by a type I specific driver, (asenseGAL4, Stinger:GFP). Compared to the mCherry control, and defining differentially expressed genes (DEGs) as those that exhibited a log2 fold change >1 and False Discovery Rate (FDR) <0.05, we identified 2,776 DEGs in the TBP RNAi (Fig 5A, left; Table S2), 152 upon TRF2 knockdown (Fig 5A, center; Table S3) and 1,165 in the TAF9 RNAi (Fig 5A, right; Table S4). Because core promoter factors are required for transcriptional activation, we focused on the down-regulated genes. Unexpectedly, most of the 1488 TBP-dependent genes were private (i.e. not co-regulated by TRF2 or TAF9), but 119 of these also overlapped with 587 genes down-regulated upon TAF9 knockdown and this overlap was statistically significant (p = 8.98 e-23, hypergeometric test; Fig 5B). Of the 94 down-regulated genes in the TRF2 RNAi condition, 45 also required TAF9 for their expression (p = 6.08 e-60 hypergeometric test). Gene Ontology (GO) analysis revealed that TBP-dependent genes were enriched for several mRNA metabolic processes and cell cycle regulation, whereas the TAF9 regulated genes were enriched for GO terms related to regulation of gene expression and transcription, as well as stem cell differentiation (Fig 5 E; Table S5, 6). In contrast, the TRF2-dependent genes did not present enriched GO categories after correcting for multiple hypothesis testing. Strikingly, the majority of TAF9-dependent genes were also private (i.e. not co-regulated by TRF2 or TBP), and included known NSC polarity genes (*insc*, *numb*, *pun*), NSC transcription factors (*HmgD*, *klu*, *chinmo, kni*) and the neuronal marker (*elav*).

**Figure 5.**
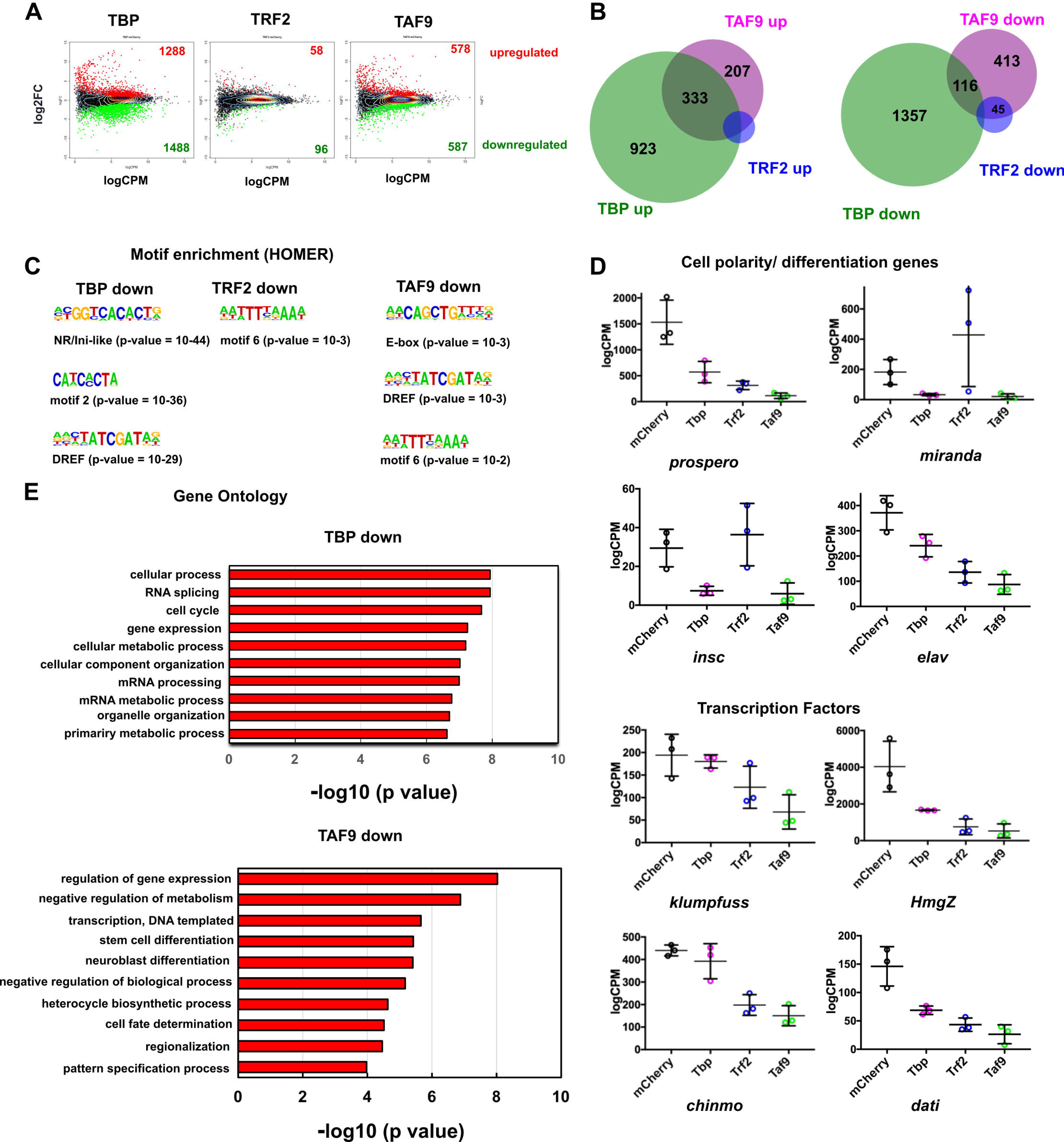
Identification of TBP, TRF2 and NSC-TAF target genes. (A) MA-plots of RNA-seq analysis and identification of differentially expressed genes (DEGs) The number of upregulated genes and downregulated genes are shown in red and green, respectively. (B) Venn diagrams show the overlap between upregulated genes (left) or downregulated (right) DEGs identified for TBP (green), TRF2 (blue) and TAF9 (magenta). (C) Motif analysis of downregulated genes using HOMER. (D) Normalized read counts of selected target genes. (E) Selected gene ontology categories that are overrepresented in gene sets downregulated upon TBP (top) or TAF9 knockdown (bottom).

For simplicity, we refer to the TBP-TAF9 co-regulated genes as NSC-TAF-TBP-dependent genes. Intriguingly, amongst the NSC-TAF-TBP targets were genes known to be involved in NSC fate determination and asymmetric cell division (*mira*, *brat*, *pon, pros*), cell cycle genes (*E2F1*, *CycE*, *stg*; Fig S5C), regulators of NSC temporal identity (*Syp*, *svp*) as well as several transcription factors (*sqz, nab, foxo, Mnt, tap*) and genes encoding RNA-binding proteins (*Rsf1, Hrb87F, B52, U4-U6-60K*; Figs 5D, S5D). We refer to genes that required both TRF2 and TAF9 for their expression as NSC-TAF-TRF2 targets. Many of these genes also encode transcription factors (*dati*, *hng3*, *HmgZ*, *mamo*, *Hr4*, *bi*, *jim*) and RNA-binding proteins (*shep*, *bru3*, *aub*, *pum*). We next performed an unbiased motif discovery of cis-regulatory sequences enriched in the TBP, TRF2 and TAF9 target genes using HOMER and unexpectedly, we found that all three sets of target genes were enriched for E-boxes, the DNA replication-related element (DRE) as well as some orphan motifs (Fig 5C). Taken together, our RNA-seq analysis revealed a complex regulatory network orchestrated by different sets of core promoter factors, with NSC-TAF-TBP regulated genes involved in cell cycle progression, NSC polarity and fate determination, whereas NSC-TAF-TRF2 targets encode a set of transcription factors and RNA-binding proteins distinct from the NSC-TAF-TBP targets.

### Identification of TBP, TRF2S and TAF5 genomic binding sites

To test whether NSC-TAFs and TRF2 co-occupy a subset of target genes that are distinct from TFIID-regulated genes identified in our RNA-seq experiments, we profiled the genomic binding sites of TBP, TRF2 and a representative NSC-TAF (TAF5) using the targeted DamID (TaDa) system. Of note, TRF2 encodes two proteins: a short isoform of 632 amino acids (TRF2S) and a longer isoform (TRF2L) of 1715 amino acids, wherein the C-terminal 632 amino acids of TRF2L are identical to TRF2S. TaDa permits spatiotemporal controlled expression of proteins of interest fused to an *E. coli* DNA adenine methyltransferase (Dam) and tissue-specific genomic profiling without cell isolation, antibodies or fixation (46). Using this system, we drove expression of UAS-Dam, UAS-Dam-TBP, UAS-Dam-TRF2S and UAS-Dam-TAF5 using a worniuGAL4, UAS-GFP, tubGAL80^ts^ driver and a 72h induction period. Genomic DNA was extracted from biological duplicates, digested with a methylation-sensitive enzyme (DpnI), amplified and the methylated GATC sites identified by next generation sequencing. Using the parameters described in the Materials and Methods section, we identified 3100 peaks for Dam-TBP, 3028 for Dam-TRF2S and 3659 for Dam-TAF5. As expected, these peaks were significantly overrepresented at and around transcription start sites (TSS; Fig 6 A, B). Analysis of overrepresented motifs surrounding TSS-associated peaks with HOMER revealed that DRE motifs, E-boxes and an Initiator-like sequence were enriched in all three Dam fusion proteins (Fig. 6D). To determine which genes are co-bound by the three Dam fusion proteins, we selected all peaks that were annotated as promoter/TSS and asked to what extent they overlap. As shown in the Venn diagram in Fig 6C, a significant number of bound genes are shared between the three Dam fusion proteins. Importantly, analysis of peaks identified in both replicates produced a similar picture (Fig S6). Consistent with our RNA-seq analysis, 955 of the TSS-associated peaks were shared between Dam-TBP and Dam-TAF5, whereas 506 peaks were shared between Dam-TAF5 and Dam-TRF2S. Moreover, this analysis also showed that Dam-TAF5 and Dam-TBP co-bound to the promoter/TSS of several of NSC-TAF-TBP-dependent genes identified above, including transcription factors (*sqz, nab, foxo, Mnt, tap*), cell cycle genes (*E2F1*, *CycE* (Fig7A), *stg*,) and RNA-binding proteins (*Syp* (Fig 7B), *Rsf1, Hrb87F, B52, U4-U6-60K*). However, the Dam-TAF5 and Dam-TRF2S co-bound genes included those encoding transcription factors (*dati*; Figure 7B, *mamo*, *Hr4)* and the RNA-binding protein *bru-3* (Fig 7C). Strikingly, some of the TAF9-dependent genes that did not require TBP for their expression were nonetheless co-bound by Dam-TAF5 and Dam-TBP, including *klu* (Fig 7D) and *insc* (Fig 7E). Taken together, the gene ontology and motif analysis of our RNA-seq data, combined with identification of genomic binding sites for this network of core promoter factors using DamID-seq, support the notion that NSC-TAF complexes directly regulate multiple aspects of NSC identity, including NSC cell polarity, cell cycle progression and RNA metabolism.

**Figure 6.**
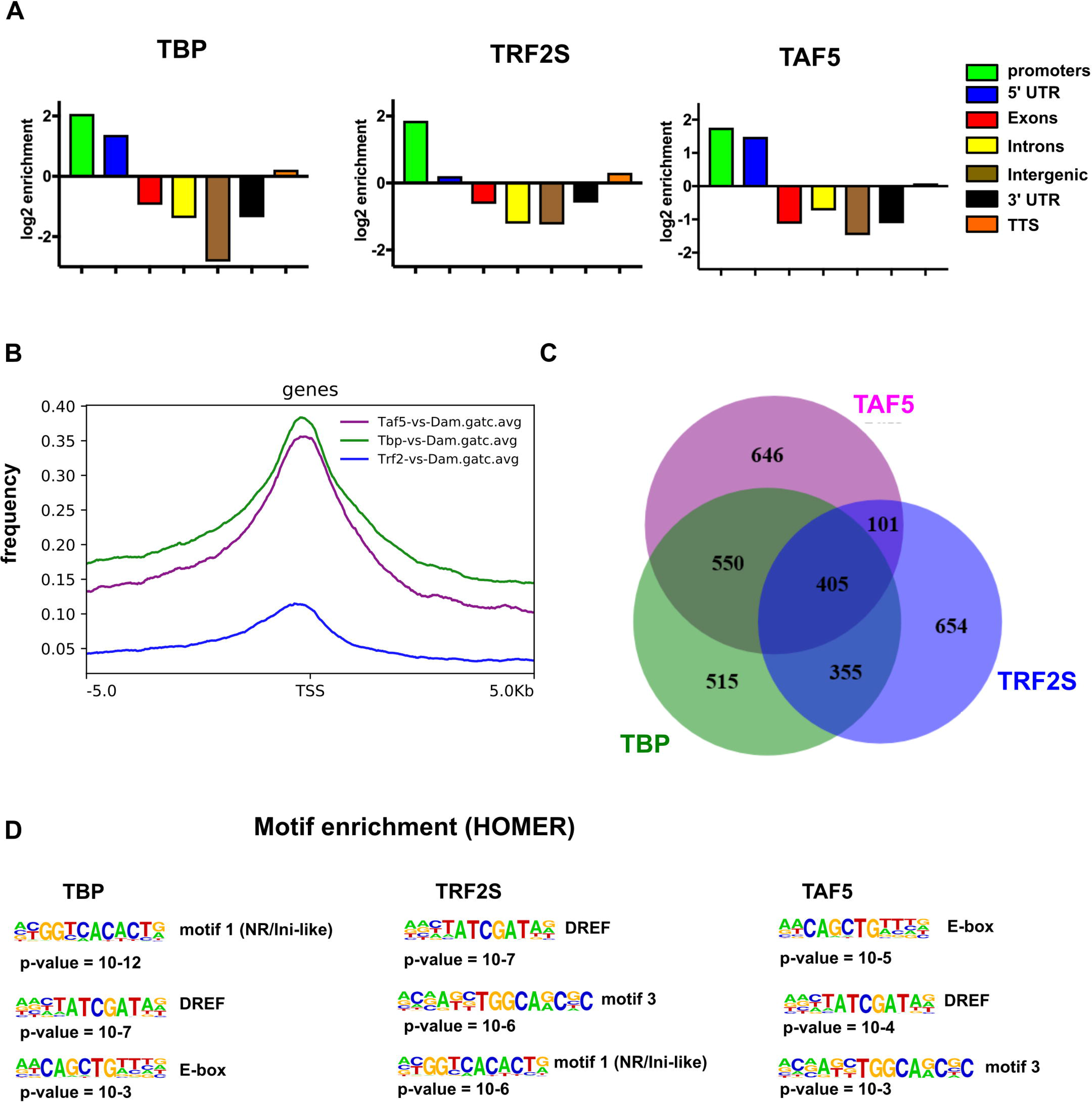
Identification of bound genomic regions by DamID-seq. (A) Genomic distribution of regions bound by Dam-TBP, Dam-TRF2S and Dam-TAF5 shows enriched binding at promoters and 5’UTRs. Plots show log2 enrichment scores (observed/expected). (B) Metagene profile of DamID-seq data for Dam-TBP (green), Dam-TRF2S (blue) and Dam-TAF5 (magenta). (C) Venn diagram showing the statistically significant overlap between genes with TSS-associated peaks. (D) Motif enrichment analysis of peaks identified motifs similar to those identified in the RNA-seq analysis.

**Figure 7.**
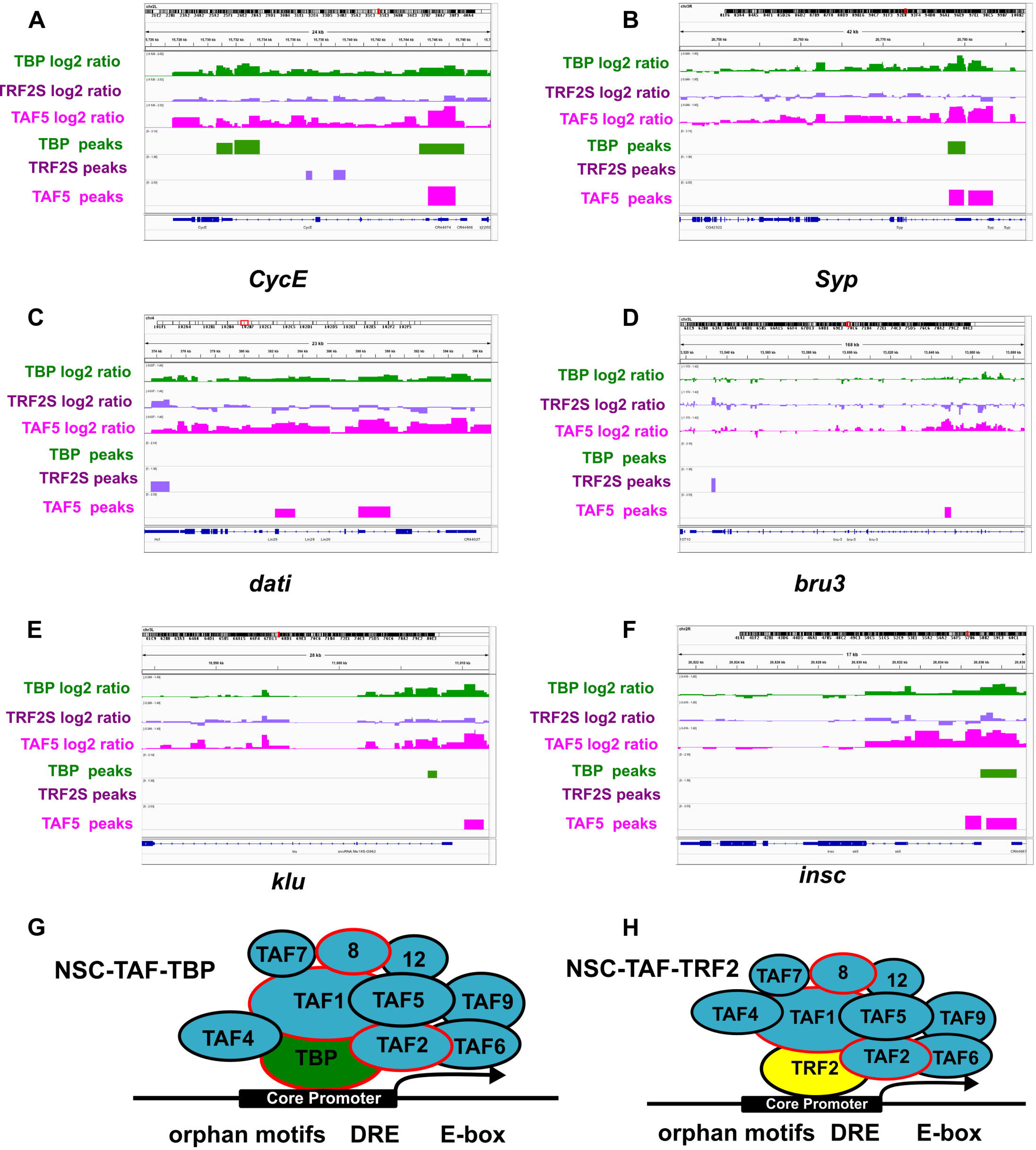
Model for regulation of TBP, TRF2 and NSC-TAF target genes. (A-F) Screenshots of DamID-seq tracks showing the log2 ratios for both replicates and identified peaks for TBP (green), TRF2S (purple) and TAF5 (magenta) for *CycE* (A), *Syp* (B), *dati* (C) and *bru3* (D), *klu* (E) and *insc*; (F) A putative NSC-TAF-TBP complex regulates a subset of NSC-expressed genes that enriched for DREs, E-boxes and orphan motifs. (G) A putative NSC-TAF-TRF2 complex regulates a different subset of NSC-expressed genes that are also enriched for DREs, E-boxes and orphan motifs (H).

## Discussion

Control of developmental gene expression is thought to primarily rely on the combinatorial action of sequence-specific transcriptional activators and repressors. The concerted action of activators and repressors in turn, is thought to ultimately converge on a highly conserved general transcription factor, TFIID. Based on studies in yeast and cultured mammalian cells, TFIID, composed of TBP and 13 TAFs, has been historically considered to be an essential yet passive player in gene regulation. However, the emergence in metazoans of both TBP and TAF paralogs, and of additional core promoter elements, has been proposed to contribute to the evolution of bilaterians by supporting more complex transcriptional programs (33). Evidence from genetic and biochemical studies in a wide variety of model systems suggest that this diversity has indeed allowed multiple TAFs to take on cell- or tissue-specific functions. For example, the TAF9 paralog TAF9B regulates neuronal gene expression by associating with the SAGA/PCAF co-activator complex, whereas the TAF7 paralog TAF7L associates with TBP-related factor 2 (TRF2) to direct expression of a subset of post-meiotic genes during spermiogenesis (11, 47). However, these examples are generally restricted to orphan TAFs, while prototypical TAFs are primarily present in TFIID and/or SAGA complexes.

In this study, using *Drosophila* NSCs as a model, we uncovered gene-selective functions for a subset of TAFs, that we refer to as NSC-TAFs, and some of these functions are shared with TRF2 whereas others are shared with their canonical binding partner, TBP. Importantly, we provide evidence that this network of core promoter factors is not required for global gene expression in NSCs. First, depletion of NSC-TAFs, TBP or TRF2 did not impact NSC survival. Second, RNA-seq analysis revealed that only 36.2%, 0.2% and 15.2% of protein-coding genes detected by our RNA-seq experiments were affected upon knockdown of either TBP, TRF2 or TAF9, respectively, suggesting that at least when examining steady-state RNA levels, that this network is dispensable for the expression of the majority of genes expressed in NSCs. Third, NSC-TAF and TRF2 functions were also process-selective, as we found that they were required for NSC cell cycle progression and NSC cell polarity but did not appear to be critical for NSC survival. Our finding that NSC-TAFs did not regulate survival was unexpected, as deletion of *Taf9* in chicken DT40 cells, of *Taf4a* in mouse embryos or depletion of TAF9 in wing disc epithelial cells all result in increased apoptosis (38, 48, 49). However, it’s unclear whether TAFs are required for survival of embryonic stem cells (ESCs) as inducible depletion of TAF8 resulted in ESC cell death in one study whereas no cell death was detected upon knockdown of either TAF5 or TAF6 in a different study (14, 19). In contrast to a report using murine ESCs (18) in which a stable TAF5 knockdown ESC line prematurely differentiated without affecting the cell cycle, we show here that NSC-TAFs and TRF2 control stem cell identity in part, through direct regulation of the cell cycle. Depletion of NSC-TAFs or TRF2 by RNAi diminished the number of NSCs that incorporated the thymidine analog EdU, lowered the mitotic index, and rendered NSCs hypersensitive to cell cycle manipulation. Moreover, we identified DamID peaks at key cell cycle genes, including *E2F1*, *CycE* and *string*. We initially hypothesized that NSCs depleted for NSC-TAFs or TRF2 would both exhibit an extended G1 phase and be hypersensitive to manipulation of the G1/S transition. However, quantification of cell cycle phases using a FUCCI-based reporter showed that NSCs depleted for NSC-TAFs or TRF2 were in fact primarily in G2 and exhibited hypersensitivity to manipulation of both the G1/S and G2/M transitions. This is intriguing because a recent study showed that quiescent NSCs, which are known to exhibit a primary process, arrest primarily in G2 and are labeled by *tribbles* (*trbl*), which encodes a conserved pseudokinase (50, 51). These phenotypes are remarkably similar to those observed upon depletion of NSC-TAFs or TRF2 and we note that the *trbl* locus is occupied by TBP, TRF2S and TAF5.

Because NSC-TAFs and TRF2 exhibit similar loss of function phenotypes and share at least 45 target genes in addition to the type I NSC marker *ase*, we proposed that they function together to direct expression of a subset of NSC-expressed genes (Fig 7F). We tested this hypothesis by combining expression analysis using RNA-seq of FACS-purified NSCs and by determining the genomic binding sites for TBP, TRF2S and a representative NSC-TAF (TAF5), using Targeted DamID (TaDa). Our RNA-seq experiments revealed that many genes co-regulated by TBP and TAF9 are known or predicted to be important for NSC identity including the chromatin remodeler *domino* (27), the cell cycle genes *E2F1*, *CycE* (Fig 7A) and *string* and the temporal identity factors *Syp* (Fig 7B) and *svp*. However, the functional relevance of the NSC-TAF-TBP target genes remains to be determined. Similarly, TAF9-dependent genes that were unaffected upon TBP knockdown and that are bound by TAF5 are good candidates for mediating NSC-TAF’s function in self-renewal, such as the transcription factors Chinmo, Klumpfuss (44), HmgD and the polarity protein Insc (Fig 7 E). Lastly, while the function of the transcription factors (*dati*, *hng3*, *HmgZ*, *mamo*, *Hr4*, *bi*, *jim*) that are co-regulated by TRF2 and TAF9 and co-occupied by TRF2S and TAF5 are not well characterized, Dati (Fig 7B), Jim and Mamo have recently been identified as important components of gene regulatory networks uncovered in a single cell RNA-seq atlas of the adult *Drosophila* brain (52).

Because TRF2 neither binds the TATA box nor has sequence-specific DNA binding activity, how does the putative NSC-TAF-TRF2 complex recognize its target genes? Several lines of evidence suggest that DREF, which directly binds the DRE element, could be part of the DNA-targeting mechanism. First, depletion of DREF also results in fewer NSCs and smaller NSC lineages (29). Second, DREF is known to be part of the TRF2 complex (53). Third, motif analyses with HOMER revealed that the DRE was overrepresented in peaks identified by TaDa for all three fusion proteins. While a TRF2 complex larger than 500kDa has been purified from embryonic nuclear extracts, none of the identified TRF2-binding proteins were TAFs (53). However, a more recent identification of TRF2S-binding proteins from ovary lysates identified several TAFs (54), raising the possibility that NSC-TAFs and TRF2 form a complex in vivo.

While depletion of TBP did not result in loss of Ase expression, nor diminish the number of NSCs, we found that TBP was essential for NSC cell cycle progression and directly regulated expression of many cell cycle genes. Intriguingly, knockdown of the RNA Pol II subunit RpII33 resulted in a more severe cell proliferation phenotype than TBP knockdown (Fig 3A-B) but we note that a complication of the RNAi experiments is that TBP depletion reduced expression of UAS transgenes including the RNAi transgene itself whereas TRF2 or NSC-TAF depletion increased expression of UAS transgenes. However, given the lack of NSC loss in TBP-depleted brains, we were surprised to find that more than a third of NSC-expressed genes detected by our low-input RNA-seq were affected upon TBP knockdown. This is even more striking considering the fact that several NSC-TAFs (TAF1, TAF4, TAF8 and TAF9) were among the genes downregulated upon TBP depletion yet the NSC-TAF and TBP phenotypes are clearly different. Importantly, a study that sought to model the human neurological disorder SCA17 in *Drosophila* showed evidence that TBP is required for normal brain function, as removal of one copy of *Tbp* recapitulates some features of SCA17, such as impaired motility and age-dependent accumulation of vacuoles in the brain (23).

We reasoned that the subset of TAFs that we identified is unique, as it is distinct from any previously described TAF complex. For example, NSC-TAFs partially overlap with a subset of TAFs that appear to co-regulate the size and composition of lipid droplets in the *Drosophila* fat body with TRF2 (35), yet that study did not identify lipid droplet functions for TAF2, TAF7 or TAF8, which are NSC-TAFs. Similarly, in mouse ESCs TFIID was proposed to be integral to the pluripotency circuitry (14) yet depletion of two NSC-TAF orthologs (TAF7 or TAF8) did not affect ESCs identity.

Recent exome sequencing studies have produced compelling evidence that pathogenic variants in *TAF1*, *TAF2*, *TAF8* and *TAF13* are linked to intellectual disability and microcephaly (16–19). However, none of these variants have been modeled in vivo in part, due to a gap in our understanding of the function of TAFs during brain development. By demonstrating that TAFs are required for NSC cell cycle progression, NSC cell polarity, and act to prevent premature differentiation while not affecting survival, our work provides a foundation for future studies aimed at uncovering the causative variants in these disorders (55).

## Materials and Methods

### Fly stocks

Unless noted otherwise, all flies were raised on standard medium at 25 ° and expression of UAS transgenes performed at 29 °. For the focused RNAi screen, 10-15 virgin females from a tester strain (worniuGAL4, UAS-GFP, UAS-Dicer2; this study) were crossed to 5 males from individual RNAi lines obtained from either the Vienna Drosophila Resource Center or the Bloomington Stock Center (TRiP lines). List of all RNAi lines used can be found in Table S1. F1 progeny from this cross were raised at 29° and dissected nervous system were screened for changes in the intensity and/or pattern of the GFP signal. Other fly lines used were:

#### GAL4 drivers

worniu-GAL4, UAS-GFP; UAS-Dcr-2, apterousGAL4, tubGAL80^ts^ (56), UAS-Dicer2 (57); worniuGAL4, aseGAL80; UASCD8:GFP (Chris Doe, University of Oregon, HHMI), aseGAL4, Stinger:GFP and UAS-Dicer2; inscGAL4, UAS-CD8:GFP; tubGAL80^ts^ (Catarina Homem, CEDOC, Portugal), worniuGAL4, tubGAL80^ts^, UAS-GFP/CyO; UAS-flp, act5C>CD2>GAL4/TM6B (this study), worniuGAL4; UASp-GFP.E2f1.1-230, UASp-mRFP1.CycB.1-266/TM6B (this study).

#### Mutants

*Taf4*^*LL07382*^ (142-038), *Taf7*^*LL01754*^ (140-455), *Taf13*^*LL04552*^ (141-287) and *Trf2*^*G0071*^ were obtained from the Kyoto stock center. *e(y)1/Taf9*^*190*^ (Wu-Min Deng (38)), *Tbp*^*SI10*^ (Spyros Artavanis-Tsakonas (58)), *Taf11*^*M1*^*, Taf11*^*M5*^ (Qinghua Liu (30)) are published stocks sent by the corresponding authors. We recombined *Tbp*^*SI10*^ onto a FRTG13 chromosome using standard fly genetics. FRT82B, *prospero^17^/TM3,Sb,ry*, RK (Chris Doe, University of Oregon, HHMI). The *prospero*^*17*^ and *Taf7*^*LL01754*^ mutant alleles were recombined using standard fly genetics.

#### Transgenes

UAS-Dcr-2 (57), PCNA-GFP (59), UAS-string RNAi (GL00513), UAS-Dacapo, UAS-Wee1(60), UAS-TRF2S-mCherry; (61) Jean-Michel Gibert), TAF1-EGFP-FlAsH-StrepII-TEV-3xFlag (BL64451), UAS-TAF2-HA;(62) Konrad Basler, FlyORF), UAS-TAF5-HA (62) Konrad Basler, FlyORF), UAS-FLAG-TRF2S, UAS-TRF2L, UAS-TAF9 (35), Taf12:2XTY1-SGFP-V5-preTEV-BLRP-3XFLAG (VDRC 318745). The UAS-TRF2S-mCherry, UAS-TAF2-HA and UAS-TAF5-HA lines contained a 3xP3-RFP transgene in the insertion site that was removed by crossing to a Cre-expressing line (Bloomington stock number 34516).

#### MARCM

FRT19A, ywhsflp^122^, tub-Gal80; tub-GAL4, UAS-GFP/MKRS (Laura Buttitta), w, C155-GAL4, UAS-mCD8-GFP, hsflp; FRT40A, tub-GAL80 (Liqun Luo), w, C155-GAL4, UAS-mCD8-GFP, hsflp; FRTG13, tub-GAL80 (Liqun Luo), w, C155-GAL4, UAS-mCD8-GFP, hsflp;; FRT2A, tub-Gal80 (Liqun Luo), w, UAS-mCD8-GFP, hsflp;; FRT82B, tubGAL4, tub-Gal80 (Laura Buttitta).

### Clonal analysis

To generate MARCM clones, virgin females from MARCM-ready lines described above were crossed to control males and experimental males except for MARCM19A wherein virgin females from FRT19A, *e(y)1/Taf9*^*190*^ or *Trf2*^*G0071*^ were crossed to males from the MARCM-ready line for FRT19A. F1 larvae at 24 h after larval hatching (ALH) were heat-shocked for 1 hr at 37°C, allowed to recover for 24h at 18°C and further aged for 72h at 25°C before dissection.

### Immunofluorescence and confocal microscopy

The following antibodies were used: rabbit anti-GFP (1:500, Life Technologies A11122), chicken anti-GFP (1:250, Aves Labs GFP-1010), rabbit anti-DsRed (1:250, Clontech, 632496), mouse anti-HA (1:500, abcam 130275), mouse anti-FLAG (1:250, Sigma F3165), guinea pig anti-Deadpan (1:1000, James Skeath), rat anti-Deadpan (abcam 195172), rabbit anti-Asense (1:500, Yuh Nung Jan (63)), rabbit anti-Bazooka (1:500, (64) Fumio Matsuzaki, rabbit anti-Inscuteable (1:250, Fengwei Yu (65), mouse anti-Prospero (1:50, MR1A from DSHB), rat anti-Elav (1:100, 7E8A10 from DSHB), rabbit anti-aPKCζ;(1:500, Santa Cruz sc-216, discontinued), rabbit anti-Cyclin E (1:250, Santa Cruz sc-33748, discontinued), mouse anti-phospho-Histone H3 (1:1000, Cell Signaling 9706), rabbit anti-phospho Histone H3 (1:1000, Millipore 05-817), mouse anti-TAF4 (1:100, 3E12 clone, Robert Tjian, UC Berkeley (66), rabbit anti-TBP (1:500, James Kadonaga, UCSD (37)). Larval brains were dissected in either PBS or Schneider’s *Drosophila* media (21720-024, Thermo Fisher Scientific), fixed in 4% paraformaldehyde (Electron Microscopy Sciences) in PBS-T (PBS with 0.5% Triton X-100) for 20 min. After fixation, brains were washed in PBS-T for 10 min three times and blocked in PBS-T with 5% normal goat serum (005-000-121 Jackson Immunoresearch Laboratories) and 1% BSA (A30075-250.0, Research Products International) for 1h. Fixed brains were incubated with the relevant primary antibody solutions at the dilutions shown above for either 3h at RT or overnight at 4°. The following day, brains were rinsed and washed for 10min three times and then incubated in secondary antibody solution for 2h at room temperature. After incubation with Alexa Fluor secondary antibodies (1:400; Thermo Fisher Scientific), three 10min washes were performed and brains were mounted in Vectashield (Vector Laboratories; H-1200). Images were acquired on a Zeiss LSM780, processed with Fiji and figures were assembled with Inkscape 0.92.2. For NSC quantification, z-stacks of brain lobes were acquired with a step size of 2 μm and Central Brain NSCs (Large Dpn^+^ cells) were counted manually using the Cell Counter plugin in Fiji (67).

### NSCs sorting and RNA sequencing

Wandering third instar larvae from control or experimental genotypes were collected and washed once in phosphate-buffered saline (PBS), dissected in supplemented Schneider’s medium (10% fetal bovine serum, 2% Pen/Strep (15140122, Thermo Fisher Scientific), 20mM L-Glutamine (G7513, Sigma-Aldrich), L-Glutathione (G6013, Sigma-Aldrich) 5 mg/mL, Insulin (I0516, Sigma-Aldrich) 20 mg/mL, 20-Hydroxiy-Ecdysone (H5142, Sigma-Aldrich) 1 mg/mL) and brains were collected and washed in cold Rinaldini solution. Next, they were enzymatically dissociated in Rinaldini solution with 1mg/mL collagenase I (C0130, Sigma-Aldrich) and 1mg/mL of papain (P4762, Sigma-Aldrich) for 1hr at 30°C. Brains were washed twice with Rinaldini solution and twice more with supplemented Schneider’s medium. Brains were manually disrupted with a pipette tip in 200 μl supplemented Schneider’s medium. After filtering the cell suspension using a Flowmi tip strainer (H13680-0040, Bel-Art SP Scienceware), NSCs from control or experimental genotypes were sorted in triplicate from brains expressing mCherry, TBP, TRF2, TAF5 or TAF9 RNAi transgenes on a BD Aria 2 essentially according to (68), except that we used a 100μm nozzle and added DAPI (D9542, Sigma) at 1μg/μl to exclude dead cells. We obtained 10-15,000 putative NSCs from each sort. Sorted NSCs in TRIzol were homogenized with a pestle and then RNA was extracted according to the manufacturer’s instructions, using Glycoblue 500μg/μl (AM9516, Ambion) as a co-precipitant. RNA was then eluted in 15μl of RNase-free water. Next, we used SMART-seq v4 ultra-low input kit (634889, Clontech) to generate double-stranded cDNA for each replicate, using 1ng of total RNA as input. 1ng of the resulting cDNA, amplified with 11 cycles of PCR in the cDNA amplification step, was used to prepare sequencing libraries with the Nextera XT kit (Illumina). Paired-end sequencing (50 bp) was performed on an Illumina HiSeq 2500 and the quality control matrix was generated for all samples using RNA-SeQC (69). RNA-seq reads were aligned to the *Drosophila melanogaster* dm6 genome using TopHat v2.1.0 (70) Gene counts were generated using the Python package HTSeq v0.6.0 (71) with the “intersection-strict” overlap mode. Differentially expressed genes, defined as |log2 (ratio)| ≥1 with the FDR set at 5%, were identified using the Bioconductor package edgeR, v3.16.5 (72) after removing lowly expressed genes. The quality control steps described above prompted us to exclude the TAF5 samples. Overrepresented GO terms for genes that were either significantly up or downregulated were identified using the Bioconductor package goseq, v1.26.0. RNA-seq data has been deposited with the Gene Expression Omnibus under series GSE120433, subseries GSE120430.

### Targeted DamID (TaDa)

To generate TaDa transgenes, cDNAs for TBP (LD44083), TRF2S (LD27895) and TAF5 (LD42828) were cloned in frame into pUASTattB-LT3-NDam plasmid (kindly provided by Andrea Brand) with ϕC3-mediated site-specific integration at the attP2 site on chromosome III (strain genotype y w67c23; P {CaryPattP2). Transformants were generated by BestGene Inc (Chino Hills, CA). The UAS-Dam, UAS-Dam-TBP, UAS-Dam-TRF2S and UASDam-TAF5 transgenes were induced for 72h at 30° using a worniuGAL4, UAS-GFP, tubGAL80^ts^ driver. Genomic DNA from 100 nervous systems each was extracted in duplicate, processed and amplified essentially as described in (73), except that removal of DamID adaptors was performed before DNA shearing on a Covaris S2. Library preparation and 50bp single end sequencing was performed on a HiSeq2500 by the Fred Hutch genomics shared resource. Sequencing data was processed via damidseq_pipeline (74) (https://owenjm.github.io/damidseq_pipeline/). This pipeline includes steps of sequence alignment, read extension, binned counts, normalization, pseudocount addition and final ratio file generation. Raw reads were mapped to the *Drosophila melanogaster* genome (ftp://ftp.flybase.net/genomes/Drosophila_melanogaster/dmel_r6.14_FB2017_01, version dmel_r6.14_FB2017_01). The find peaks software (https://github.com/owenjm/find_peaks) was used to call significant peaks present in the dataset. The output log2 ratio files was converted to TDF for viewing in IGV using igvtools. The annotation and further analysis of significant peaks were done using HOMER (http://homer.ucsd.edu/homer/index.html, version v4.9). Principal Component Analysis and Spearman correlation coefficients of the read counts showed high correlation between the two sequencing runs (Fig S6A, B), but because there was some variability between peaks that were called, and we wanted to identify genes co-bound either by TBP and TAF5 or between TRF2S and TAF5, we used a less stringent FDR of 0.05 and considered peaks identified in either replicate. The DamID-seq data has been deposited with the Gene Expression Omnibus under series GSE120433, subseries GSE120432.

## Acknowledgements

We thank Spyros Artavanis-Tsakonas, Andrea Brand, Laura Buttitta, Shelagh Campbell, Wu-Min Deng, Chris Doe, Jean-Michel Gibert, Catarina Homem, Yuh Nung Jan, James Kadonaga, Qinghua Liu, Liqun Luo, Fumio Matsuzaki, James Skeath, Robert Tjian and Fengwei Yu for providing flies and/or antibodies, the Developmental Studies Hybridoma Bank (DSHB) for antibodies, the Bloomington Stock Center (NIH P40OD018537), TRiP at Harvard Medical School (NIH/NIGMS R01-GM084947), the Kyoto Stock Center and Vienna Drosophila Resource Center (VDRC) for providing fly stocks for this study. We are also indebted to the imaging, flow cytometry and genomics shared resources staff at Fred Hutch, in particular to Ms. Qing Zhang.

